# Systematic fusion transcript discovery in mantle cell lymphoma using long-read sequencing

**DOI:** 10.64898/2026.01.16.699780

**Authors:** Lael Pasipamire, Jacob Rashid, Caiden Lukan, Nandita Das, Jie Li, Chioniso Patience Masamha

## Abstract

Fusion transcripts are composed of hybrid RNA consisting of transcripts from two distinct genes and can arise from physical linking of genes at the DNA level, splicing or read-through transcription. In addition, there are also fusion transcripts that can occur between a protein coding gene and long non-coding RNAs. Systemic detection of all fusion transcripts at the RNA-level is important in the identification of potential therapeutic drug targets as well as biomarkers for detection, classification, and subtyping of cancer. We used long-read third-generation sequencing of RNA, Iso Sequencing to identify fusion transcripts in Mantle Cell Lymphoma (MCL) cell lines. Our results revealed widespread transcript diversity in MCL. The majority of the long-read transcripts were novel. Some of the thousands of novel transcripts we identified were fusion transcripts. These fusion transcripts had some of the longest transcripts in the MCL transcriptome. We identified the fusion junction of several select fusion transcripts involving protein coding genes including the well-known and widely expressed *CTBS::GNG5* and validated their presence using other techniques. Furthermore, we also identified and validated a novel fusion transcript between the multifunctional, m6A methylation ‘writer’, *RBM15,* and *LAMTOR5:AS,* a long noncoding RNA. Use of the chemical compound, JT-607, an inhibitor of CPSF73/CPSF3 which affects both alternative polyadenylation and read-through transcription resulted in increased expression of the *RBM15::LAMTOR5:AS* fusion transcript. Our analysis suggests that *RBM15::LAMTOR5:AS* and many fusion transcripts we identified are intrachromosomal. Since the origin, significance and impact of many fusion transcripts remain unknown, our results support using an unbiased approach to identify fusion transcripts. This will help us to fully comprehend the complexity of the human transcriptome in normal biology and in disease.

## INTRODUCTION

Traditionally, fusion transcripts are products of fusion genes which arise from chromosomal structural rearrangements, including deletions, insertions, inversions and chromosomal translocation events and can also be detected at the DNA level (as reviewed^1^). Dorney *et al* suggests redefining the traditional view of fusion transcripts to include any type of RNA hybrid transcript regardless of its gene annotation or its generation ^2^. Hence, in some other cases, RNA fusion events are only detectable at the RNA level but not at the DNA level. Although they are often overlooked, these RNA based fusion transcripts account for an estimated 18% of fusion events ^3^. These include fusion transcripts that are a result of transcriptional read-through of adjacent genes, also known as transcription-induced chimeras ^4^. Some fusion transcripts are as a result of splicing (cis- or trans- splicing) ^5–7^. Regardless of the source, fusions, mostly through their protein products, drive an estimated 16.5% of cancers ^8–10^. The most common well known fusion gene is *BCR::ABL1* in chronic myeloid leukemia. The *BCR::ABL1* fusion and its fusion protein product is an aspirant model for how fusions can be used for cancer diagnosis, tumor subtyping, and prognosis ^11^. The development of the bcr-abl1 fusion protein targeting drug Imatinib provided a prototype to follow in the development of fusion molecularly targeted drugs ^11,12^. This same approach can potentially be applied to other oncogenic fusions in other cancers.

The hematologic malignancy, Mantle cell lymphoma (MCL) is an aggressive subtype of Non-Hodgkin’s Lymphoma arising from B-cell lymphocytes ^13^. Despite recent therapeutic advances and initial response to therapy, relapse is inevitable and the disease is considered largely incurable more so for patients with relapsed and refractory disease^14,15^. Better understanding of the cellular and molecular biology involved in MCL oncogenesis may help further stratify patients for treatment ^15^. Like other tumors, many critical biological characteristics of MCL are retained from the untransformed cellular progenitors ^16^. Normally, B-cell lymphocytes undergo V(D)J recombination, class switch recombination as well as somatic hypermutations which create intermediate DNA breaks that precede chromosomal translocation events which mediate antigen receptor gene diversification ^17,18^. The initiating lesion for MCL is a t(11:14) translocation event that places the cell cycle regulatory oncogene cyclin D1(*CCND1*), under the control of the more active IgH heavy chain gene enhancer region and results in its constitutive expression ^13,19^. This accidental IgH recombination event has been detected in ∼2% of healthy individuals and precedes the lymphoma diagnosis by several years ^13,19,20^. Cyclin D1 expression directly affects cyclin dependent kinase independent transcriptional activity of genes that regulate chromosomal instability in human cancer ^21,22^. Hence, chronic cyclin D1 expression results in uncontrolled cell proliferation leading to high levels of DNA instability in MCL ^23–25^. Secondary genetic alterations involving chromosomal alterations, gene mutations and alterations in genes from several different pathways all contribute to MCL disease pathology ^26,27^. Aggressive MCL has high levels of aneuploidy, and has complex chromosomal alterations including chromosomal losses, chromosomal breakage-fusion bridge cycles, chromothripsis all characteristic of its instable genome ^27^. The consequences of these chromosomal abnormalities on the transcriptome are still under investigation. Although, cytogenetic analysis have revealed complex chromosomal rearrangements in MCL, little is known about the resultant fusion transcripts ^28^, as well as fusion transcripts from other causes.

Considering their potential diagnostic, prognostic and potential for targeted therapy, it is important to be able to identify all fusion transcripts regardless of their origin. In the past, specific fusion genes were identified by several approaches including chromosome banding techniques, comparative genomic hybridization arrays, fluorescence in situ hybridization and immunohistochemistry (as reviewed ^29,30^). More recently, specific gene fusions as well as specific fusion transcripts have also been identified using different forms of polymerase chain reaction (PCR), RNAse protection assays, Northern blotting and short-read sequencing (as reviewed^29,30^). The advent of next-generation sequencing technologies has allowed for systemic identification of known and novel fusion transcripts. Short-read DNA Sequencing and RNA Seq. coupled with bioinformatics has uncovered a plethora of fusion genes which still need experimental validation ^31^. While short-read RNA Seq. is a cornerstone for transcriptome analysis, it still presents challenges in transcript assembly especially for fusion transcripts.

Amplification and fragmentation steps during library preparation produce errors and biases in the obtained short reads which may result in ambiguous assignment of fragmented short reads to gene products ^32,33^. The RNA-Seq Genome Annotation Assessment Project (RGASP) involving developers of leading RNA Seq software programs looked at 25 transcript reconstruction protocols using 14 software packages on RNA Seq. data generated as part of the ENCODE project. The automated methods were unable to identify all the constituent exons for most human transcripts ^33^. When it comes to detecting structural variations, short reads had a false positive as well as false negative rate of more than 50% ^34,35^. There are over 30 different software packages, fusion detection algorithms and tools to identify gene fusions from RNA Seq. data most of which rely on the identification of paired-end reads which overlap/span the fusion juncture ^36–39^. However, these bioinformatic tools have different sensitivity and specificity making *de novo* fusion detection from bulk RNA Seq. data a major challenge ^38,39^. The advent of long-read sequencing technologies would in theory eliminate any issues associated with transcript reassembly seen in short read RNA Seq. data and eliminate bioinformatic bias against detecting fusion transcripts found only at the RNA level ^2^,

Despite the high level of chromosomal instability, reports on chromosomal abnormalities in MCL are usually obtained from individual patient case reports and little is known about the resultant fusion transcripts ^28^ as well as fusion transcripts resulting from other causes. In this paper our goal was to systematically identify fusion transcripts in MCL, regardless of their origin. To achieve this we performed third generation, PacBio long-read, Iso-Seq on RNA derived from MCL cell lines. Using SQANTI3, we classified new novel isoforms across the MCL transcriptome. Using custom scripts we identified fusion transcripts and sequences spanning the fusion junction. Next, we developed primers that spanned the fusion transcript at the 5′ and 3′ end and validated a number of those fusion transcripts. Hence, our results confirm that long-read sequencing technologies are useful in uncovering fusion transcripts regardless of their origin and can be used to fully elucidate the human transcriptome.

## RESULTS

### Characterization of long-read data

In order to identify full length RNA transcripts in Mantle Cell Lymphoma (MCL) we used long-read third generation PacBio sequencing Iso-Seq on three MCL cell lines. The RIN numbers ranged from 6.2 to 7.7. We obtained over 3 million reads per sample. A summary of the sequencing results is shown in Table 1. Shown is a representative histogram obtained from one sample and the transcript lengths ranged from ∼1,000bp to around 8000bp (Figure 1a).

**Figure 1.**
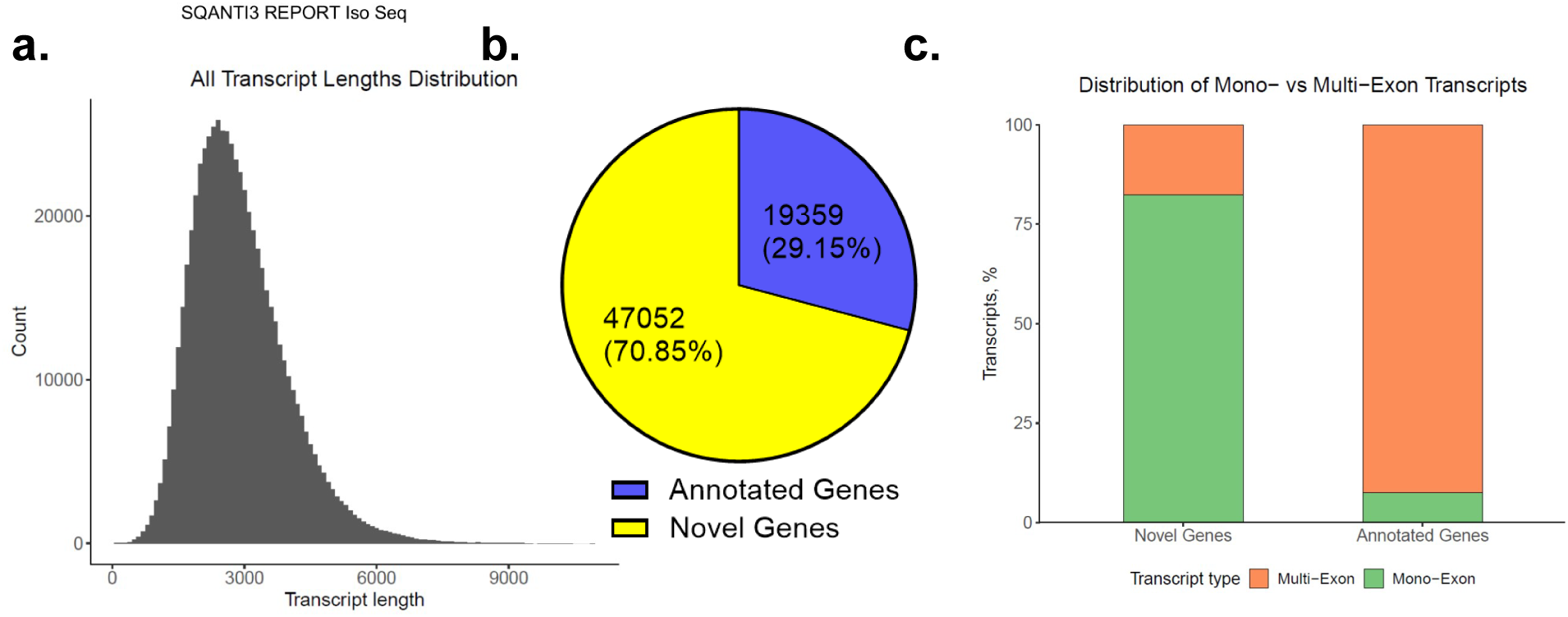
The count quantification of isoforms from Mantle Cell Lymphoma (MCL) cell lines. **a.** A representation of the transcript counts and read lengths for one sample. SQANTI3 characterization showing **b.** Annotated genes and novel genes**. c.** The percentage distribution of mono-exonic and multi-exonic transcripts belonging to novel genes and annotated genes.

**Table 1:**
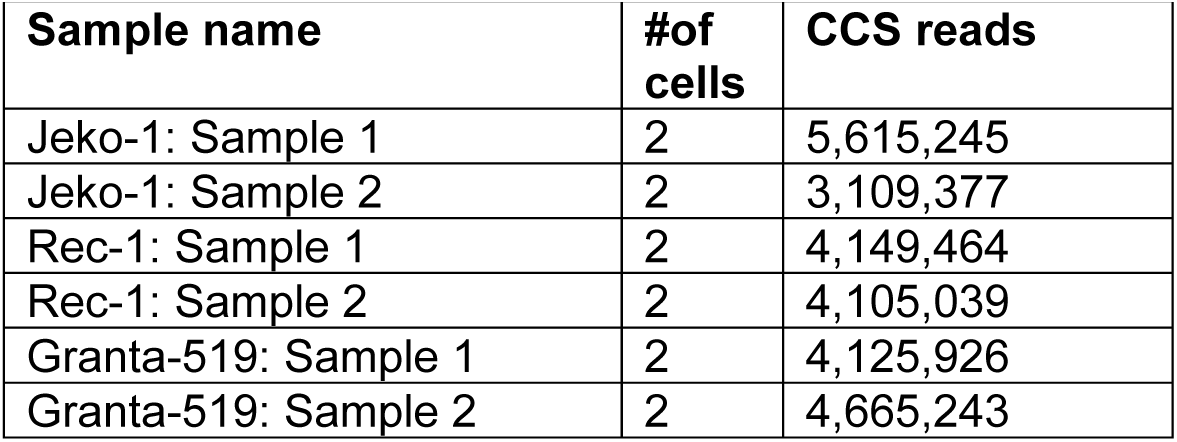
Iso Sequencing Results Summary.

### Classification of transcripts and splice junctions

A major advantage of long-read sequencing is the potential identification of novel transcripts. In order to fully annotate the Iso-Seq data, we used SQANTI3 ^40^. Interestingly, only 29.15% of our transcripts were from 19,359 known genes while the rest were categorized as novel genes (Figure 1b). The majority of the novel genes were made up of a single exon while the annotated genes consisted of multiple exons (Figure 1c). SQANTI3 can categorize each isoform based on its splice junction matches. Transcripts that match to the known reference are classified into Full Splice Matches (FSM) which completely match all splice junctions and Incomplete Splice Matches (ISM) which partially matches the reference. Novel transcripts are classified into Novel in Catalog (NIC) where the novel isoform consisting of known splice sites, Novel Not in Catalog (NNC) where novel isoforms contain at least one new splice site, genic genomic, antisense, fusion, intergenic, and genic intron ^40,41^. Our SQANTI3 classification showed that based on read counts, most transcripts were ∼2kb long and the most prevalent class of transcripts were NNC (Figure 2a). When compared by percentages, the shortest transcripts,<1kb in length, predominantly had transcripts classified as FSM followed by ISM (Figure 2b). The longest transcripts ∼10kb were predominantly classified as NNC followed by both fusion and FSM with ISM in the minority (Figure 2b). Based on read counts fusions also had relatively long read lengths (Figure 1c). Further analysis of the fusion transcripts, described by SQANTI as a transcript spanning two annotated genes, the majority of transcripts were classified as coding for multi-exonic (Figure 2d). Almost 70% of the fusion transcripts were multi-exonic and the remainder contained retained introns (Figure 2d).

**Figure 2.**
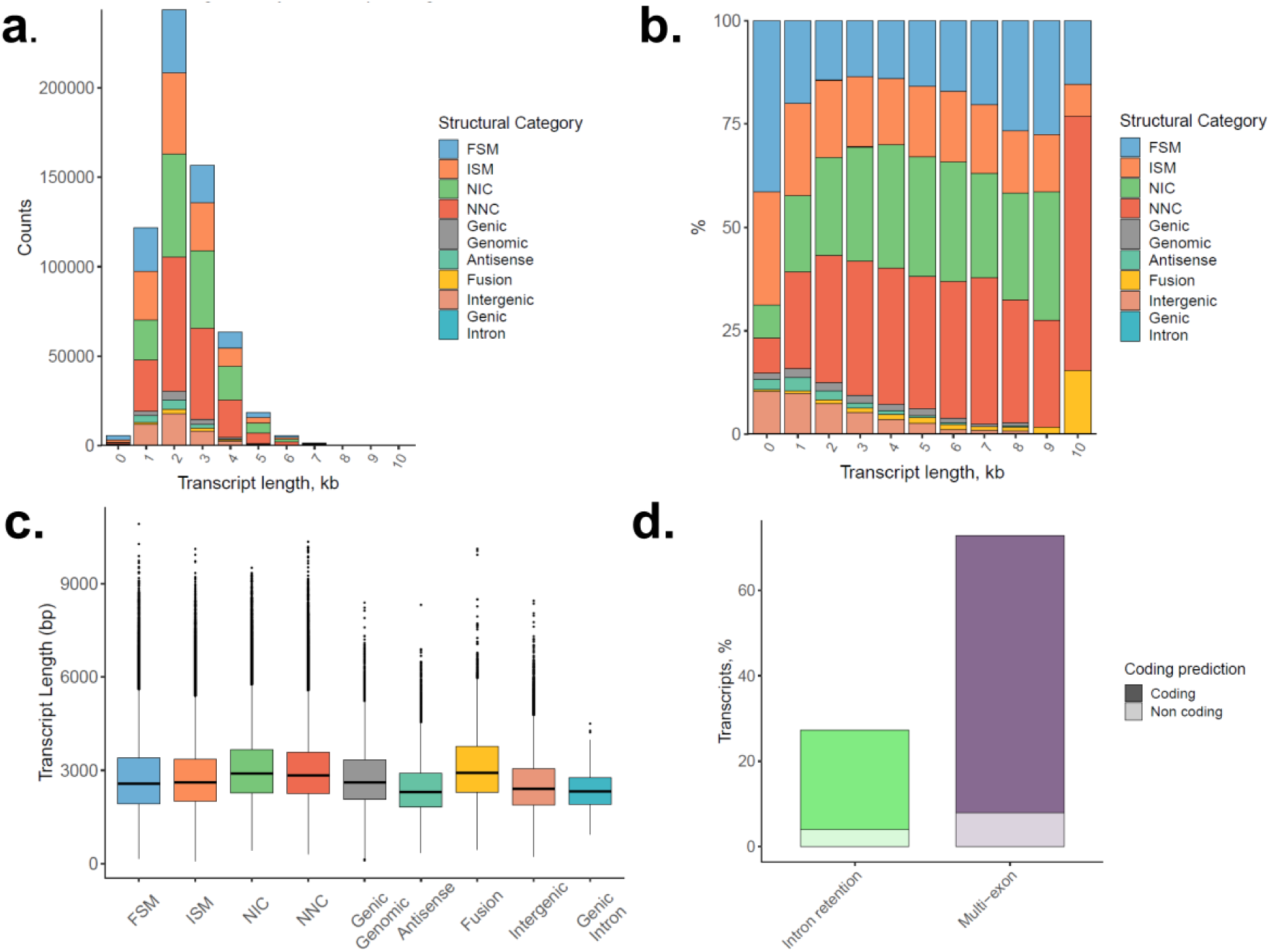
SQANTI3 classification of the MCL Iso-Seq transcripts into different structural categories. Shown are: **a**. The counts and **b**. the percentage for each structural category based on transcript length. **c**. The average transcript length for each structural category is provided. **d**. The percentage transcript distribution across different fusion categories. The structural categories include Full Splice Match (FSM), Incomplete Splice Match (ISM), Novel in Catalog (NIC), Novel Not in Catalog (NNC).

### Gene to gene fusion transcript identification and validation using PCR and Sanger Sequencing

While research suggests that there are different genomic and transcriptomic signatures that affect patient prognosis in MCL, the complete identity and role of fusion transcripts of different origins including transcriptional read-through and chromosomal aberrations are still not fully known ^42,43^. For the fusion transcripts identified in this paper we will use the recommended, double colon (::) nomenclature, regardless of the source ^44^.

After detection by RNA Seq., fusion transcripts must be carefully validated using a different but rigorous approach. FISH or PCR have commonly been used in this regard ^2^. To validate the fusion transcripts identified from the Iso-Seq data, we designed PCR primers that span the fusion breakpoint of select fusion transcripts. The forward primer was located 5’ to the fusion junction and the reverse primer was located 3’of the fusion junction. First we wanted to verify fusion transcripts occurring between two protein coding genes. As a proof of concept, we first validated a known fusion transcript, *CTBS::GNG5*, consisting of chitobiase(*CTBS*) and G protein subunit gamma 5 (*GNG5*). *CTBS::GNG5* is expressed in several tissues and both non-cancerous and cancerous cell lines ^45,46^. We detected the *CTBS::GNG5* fusion transcripts in two MCL cell lines, Jeko-1 and Granta-519 (Figure 3a), as well as in all five MCL patient derived samples (Figure 3b). We also validated the sequences at the fusion junction (Figure 3a). The *CTBS::GNG5* fusion junction was between exons and is expected to have an impact on the makeup of the protein (Table 2).

**Figure 3.**
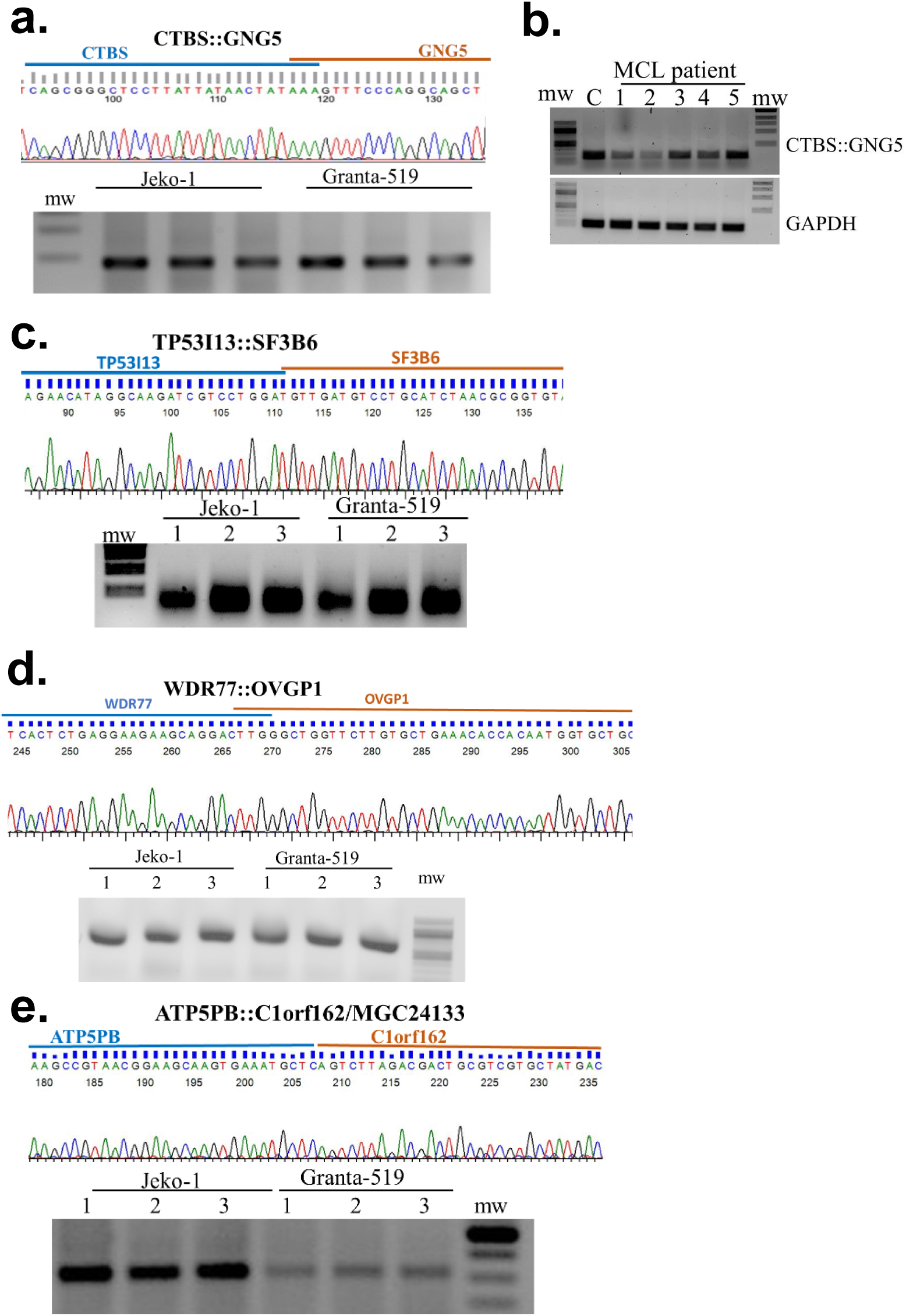
Verification of select gene to gene fusion transcripts involving coding genes in mantle cell lymphoma (MCL). **a**. The Sanger Sequencing(top panel) of the *CTBS::GNG5* PCR products extracted from the ethidium bromide stained agarose gel from two cell lines. **b**. An ethidium bromide stained agarose gel of CTBS::GNG5 PCR products from RNA derived from MCL patients and a control B-cell 19+ cell line. The top panel shows the Sanger Sequence, and the bottom panel shows the ethidium bromide gel of PCR products from two cell lines for the **c.** *TP53I13::SF3B6* **d.** *WDR711::OVGP1* and **e.** *ATP5FB::C1orf162* fusion transcripts.

**Table 2:**
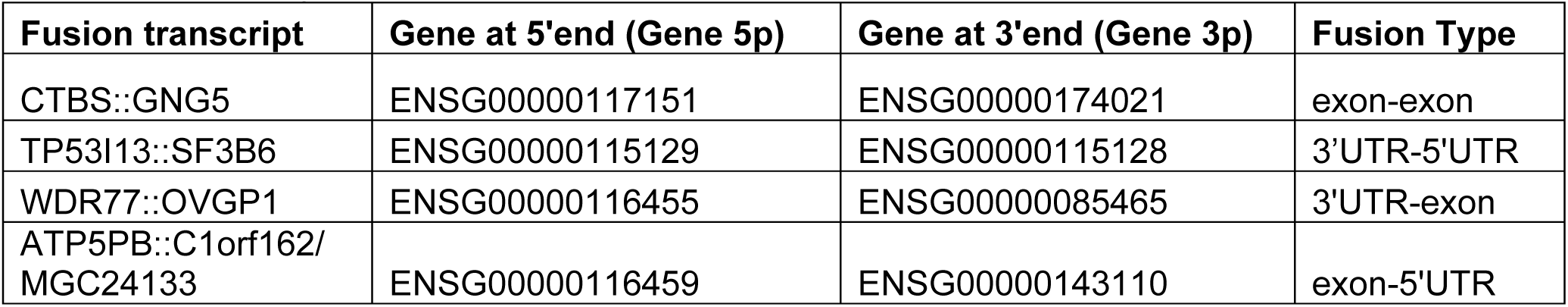
Summary of select fusion transcripts.

| **Fusion transcript** | **Gene at 5'end (Gene 5p)** | **Gene at 3'end (Gene 3p)** | **Fusion Type** |
| --- | --- | --- | --- |
| CTBS::GNG5 | ENSG00000117151 | ENSG00000174021 | exon-exon |
| TP53I13::SF3B6 | ENSG00000115129 | ENSG00000115128 | 3'UTR-5'UTR |
| WDR77::OVGP1 | ENSG00000116455 | ENSG00000085465 | 3'UTR-exon |
| ATP5PB::C1orf162/<br>MGC24133 | ENSG00000116459 | ENSG00000143110 | exon-5'UTR |

Next we validated the presence of the fusion transcript consisting of sequences from the tumor protein p53 inducible protein 3 (*TP53I13*) and splicing factor 3b subunit 6 (*SF3B6*)/Pre-mRNA branch site protein p14 (*SF3B14a*). We detected the *TP53I13::SF3B6* fusion transcript in both Jeko-1 and Granta-519 cell lines (Figure 3c). The *TP53I13::SF3B6* fusion was not expected to affect the SF3B6 protein (Table 2). Then we validated the presence of the fusion transcript, *ATP5FB::C1orf162,* between the ATP synthase peripheral stalk-membrane subunit b (*ATP5FB/ ATP5FB1*) and chromosome 1 open reading frame 162 (*C1orf162*)/ *MGC24133* (Figure 3d) in both cell lines. The fusion junction was 6 nucleotides upstream of the stop codon in the open reading frame of *ATP5PB* and the 5’UTR of *C1orf162* (Table 2). The last gene to gene fusion transcript we validated was between the WD repeat domain 77 (*WDR77/MEP50/p44/Nbla10071,RP11-552M11.3*) and the oviductal glycoprotein 1 (*OVGP1/CTIT5/ECP/MUC9/OGP*). *WDR77::OVGP1,* was detected in both cell lines and appears to be expressed at lower levels in Granta-519 cells (Figure 3e). The *WDR77::OVGP1* fusion transcript junction was between the 3’UTR of *WDR77* and an exon of *OVGP1* (Table 2). A comprehensive list of the fusion transcripts we detected from the long-read sequencing data is provided (Supplementary Table 1). It includes another known fusion, *PRIM1::NACA*, which is considered to be oncogenic. This *PRIM1::NACA* fusion was described as being between exons of both genes ^47,48^.

### Fusion transcripts between coding and non-coding transcripts

Some of the fusion transcripts we identified were between a coding gene and long non-coding(lncRNA) transcripts (Supplementary Table 1). One such fusion transcript we identified was between RNA binding motif protein 15 (*RBM15/OTT1/OTT*) and *LAMTOR5* antisense RNA1 (*LAMTOR5-AS/SLC16A4* antisense RNA 1 (*SLC16A4-AS1*). *LAMTOR5-AS* is a the long-non-coding RNA (lncRNA) that has been reported to regulate the *LAMTOR5/ HBXIP/XIP* gene by acting as a miRNA molecular sponge ^49^. Translation of *RBM15* was previously reported to be regulated by a different antisense lncRNA, *AS-RBM15/RBM15-AS,* which is located upstream of the 5’UTR of *RBM15* and is transcribed in the opposite direction (Figure 4a) ^50^. Our sequencing (Supplementary Table 1) showed the sequences at 5’end and the 3’end of the fusion junction (Figure 4b) which map to *RBM15* and *LAMTOR5-AS* respectively. Our analysis found that the fusion junction was located where the *RBM15* sequence ends in the 3’UTR after 3’end cleavage and *LAMTOR5-AS* starts at the first nucleotide (Figure 4a). The RBM15 portion contains the canonical polyadenylation signal, AATAAA, which would normally be used for proper 3’end formation and generation of a ploy(A) tail. The orientation of the fusion transcript showed that there would be no effect on the protein structure of *RBM15*. To verify the presence of *RBM15* we used primers that target *RBM15* alone and were able to detect *RBM15* transcripts using PCR(Figure 4c). We also developed forward primers that target the *RBM15* sequence and reverse primers that target *LAMTOR5-AS*. We used two different primer pairs for the latter and were able to detect and amplify the *RBM15::LAMTOR5-AS* (Figure 4c). When we used western blot analysis we detected the *RBM15* protein in the mantle cell lymphoma (MCL) cell lines (Figure 4d) and other cell lines (Figure 4e).

**Figure 4.**
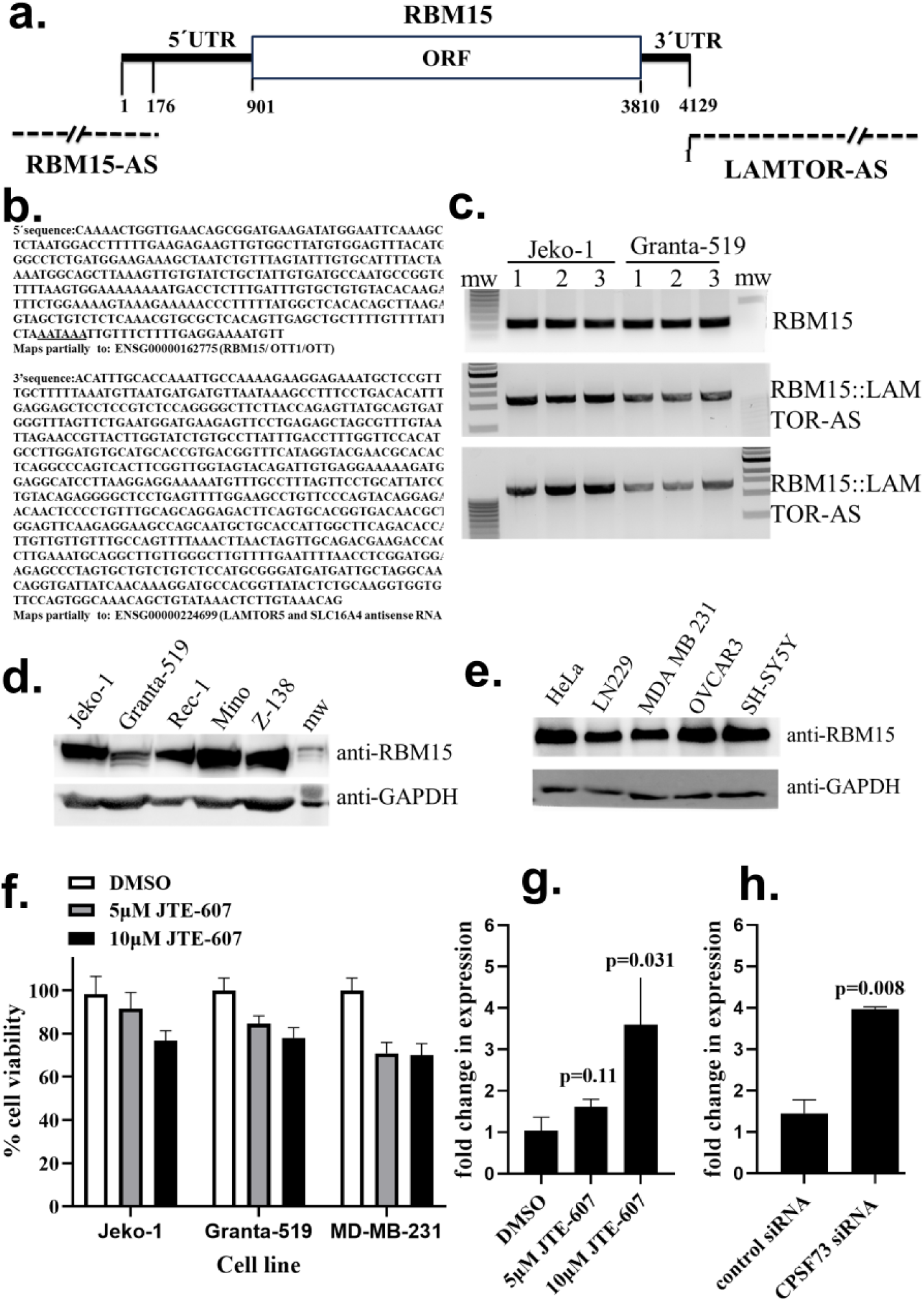
The *RBM15::LAMTOR* fusion transcript. **a**. *RBM15* Schematic representation of the human RBM15 transcript and the relative locations of the antisense *RBM15-AS* and *LAMTOR5-AS* transcripts. **b**. Iso-Seq. reads showing the sequence at the 5’end and the sequence at the 3’end of the fusion mapping to *RBM15* and *LAMTOR5-AS* respectively. Underlined is a potential polyadenylation signal within RBM15. **c.** Ethidium Bromide Agarose gel showing the PCR products obtained using primers in the ORF of *RBM15* (top panel) and two different sets of primers (set one and set 2) spanning the *RBM15* and the *LAMTOR5-AS* regions. Results of western blot analysis of cell lysates from **d**. mantle cell lymphoma cell lines and **e**. other cell lines using antibodies targeting *RBM15* with *GAPDH* as the loading control. **f**. MTT assay results for different cell lines treated with delivery vehicle (DMSO) or JTE-607 for 72 hours normalized to DMSO control (n=6, shown is the mean and SD). MDA-MB-231 qRT-PCR results showing fold change in expression of the *RBM15::LAMTOR5* fusion transcript after **g**. 72 hours treatment with JTE-607 over DMSO treated samples after normalizing to GAPDH and **h**. 48 hours after siRNA with CPSF73 over control siRNA. For **g** and **h**, shown is the mean and SD (n=3), with p-values calculated from a two tailed t-test).

Based on our analysis we posited that *RBM15::LAMTOR5-AS* is a fusion transcript resulting from read-through transcription. We decided to test the impact of the molecule, JTE-607, which works by inhibiting CPSF-73/CPSF3. One of the effects of JTE-607 is the facilitation of read-through transcription ^51,52^. Treatment of cells with 10 µM JTE-607 resulted in a decrease in cell viability for all three cell lines we tested (Figure 4f). Next we treated the MDA-MB-231 cell line, one of the cell lines that expressed lower levels of *RBM15* than other cell lines (Figure 4e) and used primers that target *RBM15::LAMTOR5-AS for qRT-PCR.* We found that treatment with 10 µM JTE-607 resulted in a significant increase (p<0.05) in the levels of the fusion transcript (Figure 4g). We also saw a significant increase in the expression of the *RBM15::LAMTOR5-AS* when we knocked out CPSF73 using RNAi (Figure 4g) in the same cell line. This suggests that inhibiting CPSF73 results in increased read-through transcription generating more *RBM15::LAMTOR5-AS* fusion transcripts.

## METHODS

### Cell culture

With the exception of the Granta-519 cell line, all the mantle cell lymphoma cell lines (Jeko-1, Mino, Z-138 and Rec-1) and other cell lines, Hela, LN229, MDA-MB-231, Ovcar-3 and SH-SY5Y) and CD19+ B-cells were purchased from ATCC (Manassas, VA). The Granta-519 cell line was purchased from the DSMZ-German Collection of Microorganisms (DSMZ, Braunschweig, Germany). The B-cells were cultured in RPMI-1640 media (ATCC, Manassas, VA) and the rest of the cell lines were cultured in DMEM with GlutaMAX (ThermoFisher Scientific, CA). In each case, cell culture media were supplemented with both 10% FBS (ThermoFisher Scientific, CA) and 1% penicillin/streptomycin (ThermoFisher Scientific, CA) at 37°C in a humidified incubator(5% CO_2_).

### MTT Assay

Cells were plated in 96 well plates (7,500 or 10,000 cells/well based on cell size). After 24 hours, cells were treated with DMSO or JTE-607 for 72 hours. Cell viability was determined carried using the MTT Cell Proliferation Assay (ATCC, Manassas, VA). In brief, 10ul of the MTT reagent was added to each well and the plate was incubated for 2 hours. Then 100ul of detergent reagent was added and the plate was incubated overnight at RT and absorbance was read at 570nm.

### RNA Extraction

After TRIzol extraction (ThermoFisher Scientific, CA), as per manufacturer’s protocol, the 260/280 values as well as the total RNA concentration were measured using a Nanodrop. To determine RNA integrity, RNA samples with 260/280 values between 2.0 and 2.10 were then run on an agarose gel. High quality RNA was then used for downstream applications.

### Iso-Seq

Iso Sequencing was done for two biological replicate samples of Jeko-1, Rec-1 and Granta-519. High quality RNA was submitted to Arizona Genomic Institute (AGI, Arizona) for further processing and Iso Sequencing using the Iso Seq Express Template PACBIO standard protocols for Single Molecule, Real-Time (SMRT®) sequencing and sequenced using the Sequel® II system. In brief, an Agilent 2100 Bioanalyzer was used for further RNA quality control and only RNA with RIN values between 6.2 and 7.7 and acceptable Bioanalyzer RNA traces were used for cDNA synthesis using the New England Biolabs NEBNext® Single Cell/Low Input cDNA Synthesis & Amplification Module (New England Biolabs, MA) with 12 cycles of amplification. Promega Pronex beads were used to purify the cDNA, and a sequencing library was prepared using Pacific Biosciences SMRTbell Express Template Prep Kit 2.0 (PacBio, CA) and sequenced on the PacBio Sequel II system using CCS mode for 30 hours.

Demultiplexed HiFi PacBio Iso-Seq data first went through the Iso-Seq pipeline (v3.4.0) to remove primer sequences, poly(A) tails and artificial concatemers. The cleaned data from all samples were merged, followed by clustering. This generated 828682 high quality transcript clusters and 460 low quality clusters. The high quality transcripts were mapped to human reference genome (GRCh38) using pbmm2 with preset parameters for Iso-Seq data. Unique isoforms were created using Iso-Seq collapse command. The gene structure gff file was input to SQANTI3 for to generate classification. The isoforms that were identified by SQANTI3 as fusion products were then analyzed using custom scripts to identify the fusion site by finding the mapping between the fusion transcript sequence to the sequences of the genes that are involved in the fusion event.

### PCR, TOPO cloning and Sanger Sequencing

High quality total RNA was converted to cDNA using RevertAid First Strand cDNA Synthesis Kit (ThermoFisher Scientific, CA). PCR was performed using Hot Start Taq 2X Master Mix (New England Biolabs, MA). For PCR, 1ul of Forward primer and 1ul of Reverse primer were added to 2ul of cDNA and 6ul of water. Then 10ul of Hot Start Taq 2X Master Mix was added and PCR was cycled following the manufacturers protocol. The PCR primers used for the *CTBS::GNG5* fusion transcript were *CTBS* Forward TGCAAAAGTCCCTTTCCGGG and *GNG5* Reverse AGATACTCCAGTCAGCAGAGGG. For the *TP53I3::SF3B6* fusion transcript the primers used were *TP53I3* Forward ATTCTGCCTCACTTCTCCACG and *SF3B6* Reverse GGGTTACACCGCGTTAGATG. To detect the *ATP5PB::C1orf162* fusion transcript the primers were *ATP5PB* Forward GCATTGCGGACCTAAAGCTG, *C1orf162* Reverse GTGTCGGGTTTACATGTGGAGCC. The primers for *WDR77::OVGP1* fusion transcripts were *WDR77* Forward GACTTGGTCCCCGCTCAATC, *OVGP1* Reverse GGTCATGGGGCAAGATCGAG. For cloning, the same primers were used to amplify the fusion transcript together with 2XPhusion Master Mix (ThermoFisher Scientific, CA) as per manufacturers protocol. After PCR, the PCR products were run on a 1% agarose gel containing Ethidium Bromide and the PCR bands were purified using the Wizard® SV Gel and PCR Clean-Up System (Promega, WI). The Zero Blunt TOPO PCR cloning kit was used to clone the PCR products as per manufacturers protocol and plasmids with inserts were sent for Sanger Sequencing at Lone Star labs (LS Labs, TX).

The primers used for Hot Start Taq 2X Master Mix (New England Biolabs, MA) PCR for *RBM15* were Forward TAAAGCTACACCCACCACCCG and Reverse CTCTAAGGCGTCGATCTGGG. The first set of primers for the *RBM15::LAMTOR5-AS* PCR were *RBM15* Forward TCCCACCTTGTGAGTTCTCCCAG and *LAMTO5-AS* Reverse GTTGTCACCGTGCACTGAAGTCTCC, and the second set were *RBM15* Forward CTGGTTGAACAGCGGATGAAGATA and *LAMTOR5-AS* Reverse TGTTCTCCTGTACTGGGAACAGGCT. For *RBM15::LAMTOR5-AS* fusion transcript we did not get any products when we used the 2XPhusion Master Mix (Thermo Fisher Scientific, CA) so we used Hot Start Taq 2X Master Mix (New England Biolabs, MA) with 25ul of the 2X master mix instead. Some of the PCR product was gel purified and used for Sanger sequencing, while the remainder was subject to T4 DNA ligation using T4 DNA ligase (New England Biolabs, MA), and the clones sent for Sanger Sequencing.

For qRT-PCR of the *RBM15::LAMTOR5-AS* fusion transcript we used the *RBM15* Forward primer GTGATGCCAATGCCGGTGTTTTAAG and *LAMTOR5-AS* Reverse primer GAACCGTTACTTGGTATCTGTGCCTTA with KAPA SYBR FAST qPCR kits (Roche, IN).

### Western Blot

Cell lysates were obtained after two freeze and thawing cycles in RIPA Lysis and Extraction Buffer (Thermo Fisher Scientific, CA) containing protease inhibitors. After quantification using Pierce™ Detergent Compatible Bradford Assay Kit (Thermo Fisher Scientific, CA), an equal amount of protein was loaded onto an 8% SDS-PAGE gel and transferred to a PVDF membrane (BioRad, Hercules, CA). After blocking in 5% milk in TBST-T buffer, cells were incubated with primary antibodies for RBM15 (ProteinTech, Rosemont, IL) at 1:1000 and GAPDH (Abcam, Waltham, MA) at 1:5000. After washes with TBS-T the membranes they were incubated with secondary antibodies, i.e. 1:5000 of anti-mouse HRP and anti-rabbit Alexa Fluor 680 Alexa Fluor™ 680 antibody (Thermo Fisher Scientific, Waltham, MA) respectively.

## DISCUSSION

Analysis of our mantle cell lymphoma (MCL) cell lines showed that the majority of our transcripts were longer than 1000 bp and our transcripts had an average read length of 3,000 nt which is similar to the median of 2,956 nt which was previously reported for the human genome ^53^.

Surprisingly, analysis of the Iso-Seq data using SQANTI3 showed that the majority of genes identified (47,052) were novel genes. Recently long read Iso-Seq analysis of human whole blood from healthy individuals identified ∼46,000 genes, with a large number of novel transcripts showing the potential of long-read sequencing to identify novel transcripts ^54^. High numbers of novel transcripts were also identified from long read sequencing of human dorsal root ganglion and this was hypothesized to be attributable to tissue specific diverse splicing unique to the central nervous system ^55^. We posit that there are MCL as well as blood specific novel transcripts whose biological significance still remains undetermined. More work still needs to be done in larger samples to determine how widespread these novel isoforms are.

MCL has been reported to have high genomic instability, with a significant number of chromosomal aberrations which are expected to give rise to fusion transcripts ^25–28^. For an unbiased view, we used our long-read sequencing data to identify fusion transcripts regardless of the source of the fusion. As proof of concept, we were able to identify the *CTBS::GNG5* fusion transcript in all our samples and validated the results using PCR in MCL cell lines and patient derived RNA. The *CTBS::GNG5* transcript generates an in frame chimeric protein and is known to be widely expressed in many tissues and cell lines ^46^. RNAi targeted depletion of *CTBS::GNG5* in the RWPE cell line, resulted in decreased cell growth and a reduction in cell motility ^46^. We also identified and validated the presence of several other fusion transcripts that would potentially impact the protein product produced at the 5’end of the fusion. These included the *TP53I13::SF3B6* and *ATP5PB::C1orf162* fusion transcripts. The *WDR77::OVGP1* fusion transcript was within the 3’UTR of the gene at the 5’end and 5’UTR of *OVGP1* which is not expected to have an impact on the makeup of the protein products.

The *RBM15::LAMTOR5-AS* fusion transcript junction is between the 3’UTR of *RBM15* and the lncRNA, *LAMTOR5* located immediately downstream of the *RBM15* gene. For *RBM15*, a previous study identified a lncRNA, *AS-RBM15* which is transcribed in the opposite direction of *RBM15* and partially overlaps with a portion of *RBM15*’s 5’UTR ^50^. The antisense, *AS-RBM15* lncRNA, as well as *RBM15* were activated by the same transcription factor RUNX1 with *AS-RBM15* increasing translation of *RBM15* with no effect on *RBM15* mRNA levels. This *AS-RBM15* appears to be distinct from the *LAMTOR5-AS* transcript we detected which is located immediately after the 3’UTR of *RBM15*. The *RBM15::LAMTOR5-AS* fusion transcript we identified is also different from another reported recurrent fusion between *RBM15 and MLK1* arising from a chromosomal translocation which results in a *RBM15::MLK1* fusion protein in acute megakaryoblastic leukemia ^56^. The impact of the fusion we discovered between *RBM15* and *LAMTOR5-AS* still remains unknown. There is no overlap in sequences between *RBM15* and *LAMTOR-AS*, and their proximity suggests that *LAMTOR5-AS* would act in cis to exert its effects (if any) on *RBM15* ^57^. Determining if the *RBM15::LAMTOR5-AS* fusion transcript affects the translation, expression, half-life or regulates *RBM15* in any way is important since *RBM15* is an RNA binding protein that has several functions including acting as one of the main drivers for N6-methyladenosine (m6A) methylation enabling m6A modification of RNAs thus regulating a large number of different processes including alternative splicing. This can potentially impact a diverse number of normal biological and disease states including cancer (as reviewed ^58,59^).

Besides *RBM15::LAMTOR5-AS,* many of the other fusion transcripts we identified from our Iso-Seq data may be intrachromosomal. The makeup of the *RBM15::LAMTOR5-AS* fusion transcript suggests that this is a read-through transcript. Read through transcripts are also known as downstream of gene containing transcripts (DoGs) ^60^. Inhibition of the endoribonuclease CPSF73/CPSF3, a component of the pre-mRNA 3’end cleavage and polyadenylation complex by JTE-607 resulted in an increase in *RBM15::LAMTOR5-AS* fusion transcript levels. This supports previous findings that inhibition of CPSF73 by JTE-607, results in an increase in read-through transcription and the generation of RNA-DNA hybrid structures ^51^. When we used RNAi to knock down CPSF73 we saw a similar increase in the levels of *RBM15::LAMTOR5-AS.* Besides *RBM15::LAMTOR5-AS,* another read-through transcript that we identified and has been previously reported from analysis of short-read RNA-Seq data is the *ATP5PB::C1orf162* fusion transcript which was identified in skeletal muscle ^61^. Thus, our results suggest that many of the fusion transcripts we identified in MCL are as a result of read-through transcription. While read-through transcription is emerging as a cellular stress response in human disease including cancer, read through transcripts are reported to be more pervasive than previously thought, occurring even in normal healthy tissues ^61,62^. The mechanisms involved in the formation of read through transcripts, as well as their biological impacts still remain largely unknown.

In conclusion, our study identified known and novel fusion transcripts and revealed the transcriptome diversity in mantle cell lymphoma. We also verified that targeting CPSF73 is a potential mechanism that may give rise to read-through transcripts.

## DATA AVAILABILITY

The Iso-Seq data that we generated for this manuscript are available for public access as of the date of publication under GEO: GSE#########.

## SUPPLEMENTARY DATA

Supplementary data is available at ######.

## AUTHOR CONTRIBUTIONS

C.P.M. developed the concept. L.P., C.L.,J.R.,C.P.M. performed the experiments. C.P.M., L.P. N.D., C.L., J.R. and J.L. analyzed the data. J.L. did the bioinformatic analysis. C.P.M. and J.L. wrote the first draft. C.P.M., L.P., N.D., C.L., J.R. and J.L. reviewed and edited the manuscript.

## FUNDING

Research reported in this publication was supported by the National Institute of General Medical Sciences of the National Institutes of Health under award number R01GM135361 awarded to C.P.M.

## Notes

### Competing Interest Statement

The authors have declared no competing interest.

